# CPPred-sORF: Coding Potential Prediction of sORF based on non-AUG

**DOI:** 10.1101/2020.03.31.017525

**Authors:** Xiaoxue Tong, Xu Hong, Juan Xie, Shiyong Liu

## Abstract

In recent years, researchers have discovered thousands of sORFs that can encode micropeptides, and more and more discoveries that non-AUG codons can be used as translation initiation sites for these micropeptides. On the basis of our previous tool CPPred, we develop CPPred-sORF by adding two features and using non-AUG as the starting codon, which makes a comprehensive evaluation of sORF. The database of CPPred-sORF are constructed by small coding RNA and lncRNA as positive and negative data, respectively. Compared to the small coding RNAs and small ncRNAs, lncRNAs and small coding RNAs are less distinguishable. This is because the longer the sequences, the easier to include open reading frames. We find that the sensitivity, specificity and MCC value of CPPred-sORF on the independent testing set can reach 88.22%, 88.84% and 0.768, respectively, which shows much better prediction performance than the other methods.

## INTRODUCTION

The so-called ncRNAs must not be coding, but it is not the truth. A large number of research reports that ncRNAs may contain small open reading frames (small ORFs, sORFs) that can encode micropeptides (Anderson, Anderson, Chang, Makarewich, Nelson, McAnally, Kasaragod, Shelton, Liou and Bassel-Duby et al. 2015; Aspden, Eyre-Walker, Phillips, Amin, Mumtaz, Brocard and Couso 2014; Bazzini, Johnstone, Christiano, Mackowiak, Obermayer, Fleming, Vejnar, Lee, Rajewsky and Walther et al. 2014; Hanada, Higuchi-Takeuchi, Okamoto, Yoshizumi, Shimizu, Nakaminami, Nishi, Ohashi, Iida and Tanaka et al. 2013; Kondo, Plaza, Zanet, Benrabah, Valenti, Hashimoto, Kobayashi, Payre and Kageyama 2010; Lauressergues, Couzigou, Clemente, Martinez, Dunand, Becard and Combier 2015; Magny, Pueyo, Pearl, Cespedes, Niven, Bishop and Couso 2013; Pauli, Norris, Valen, Chew, Gagnon, Zimmerman, Mitchell, Ma, Dubrulle and Reyon et al. 2014). The sORF encoded polypeptides (SEPs) participate in various biological processes, and they have been found in plants, bacteria, yeast, fruit flies, mice and humans, which include embryonic development (Pauli, Norris, Valen, Chew, Gagnon, Zimmerman, Mitchell, Ma, Dubrulle and Reyon et al. 2014), muscle function (Anderson, Makarewich, Anderson, Shelton, Bezprozvannaya, Bassel-Duby and Olson 2016; Anderson, Anderson, Chang, Makarewich, Nelson, McAnally, Kasaragod, Shelton, Liou and Bassel-Duby et al. 2015; Matsumoto, Pasut, Matsumoto, Yamashita, Fung, Monteleone, Saghatelian, Nakayama, Clohessy and Pandolfi 2017; Nelson, Makarewich, Anderson, Winders, Troupes, Wu, Reese, McAnally, Chen and Kavalali et al. 2016), apoptosis (D’Lima, Ma, Winkler, Chu, Loh, Corpuz, Budnik, Lykke-Andersen, Saghatelian and Slavoff 2017), prevention of polymerase II phosphorylation in germ cells (Hanyu-Nakamura, Sonobe-Nojima, Tanigawa, Lasko and Nakamura 2008), participation in cell growth and development (Casson, Chilley, Topping, Evans, Souter and Lindsey 2002; Lauressergues, Couzigou, Clemente, Martinez, Dunand, Becard and Combier 2015; Wen Lease and Walker 2004), regulation of metabolism (Lee, Zeng, Drew, Sallam, Martin-Montalvo, Wan, Kim, Mehta, Hevener and de Cabo et al. 2015) and so on. Therefore, it is great of value to make the systematic and comprehensive study of whether ncRNA can encode small peptides.

Researchers are increasingly focusing on micropeptides encoded by sORFs. In 2012, Slavoff *et al.* find 90 SEPs in human cells (Slavoff, Mitchell, Schwaid, Cabili, Ma, Levin, Karger, Budnik, Rinn and Saghatelian 2013), many of them initiated with non-AUG start codons, with non-AUG accounting for 57% and AUG accounting for 43%. This indicates that noncanonical translation is widespread in manmals. Throughout the current researches on sORF-encoded micropeptides, it is not uncommon for many micropeptides to use non-AUG as the start codon (Andrews and Rothnagel 2014; Gao, Wan, Liu, Ma, Shen and Qian 2015; Ingolia Lareau and Weissman 2011; Lee, Liu, Lee, Huang, Shen and Qian 2012; Ma, Ward, Jungreis, Slavoff, Schwaid, Neveu, Budnik, Kellis and Saghatelian 2014; Slavoff, Mitchell, Schwaid, Cabili, Ma, Levin, Karger, Budnik, Rinn and Saghatelian 2013). The non-AUG is generally near-cognate codons that differ from AUG in a single nucleotide. In 1986, Kozak *et al.* (Kozak 1986) discover that if there is no start codon AUG upstream of the RNA, the start codon is a near-cognate codon embedded in a kozak sequence box (Kozak 1986; Slavoff, Mitchell, Schwaid, Cabili, Ma, Levin, Karger, Budnik, Rinn and Saghatelian 2013). In 1989, Peabody analyzed the translation efficiency of near-cognate codons in mammals. He point out that seven of the nine near-cognate codons of dihydrofolate reductase acts as translation initiation site in mammals, and his research also find that CUG appears to be the most effective non-AUG start codon, while AAG and AGG are the lowest efficiency (Peabody 1989). In 2011, Ingolia *et al.* find that the majority of proteins in the mouse genome are translated from the classical AUG codon, but the CUG and GUG are also common as the start codons. At the same time, they also find that a large number of new sORFs exist in the mouse (Ingolia Lareau and Weissman 2011). In fact, almost all codons can be used as start codon, but the translation initiation efficiency of different codons is different (Hecht, Glasgow, Jaschke, Bawazer, Munson, Cochran, Endy and Salit 2017). In 2017, Hecht group use green fluorescent protein (GFP) and nanoluciferase to quantitatively analyze the translation initiation efficiency of all codons in E. coli (Hecht, Glasgow, Jaschke, Bawazer, Munson, Cochran, Endy and Salit 2017). They find that the AUG with the strongest translation initiation efficiency and the worst CCU are about 10,000 times different. Finally, they detect at least 47 possible start codons that can initiate protein translation. In recent years, more and more studies have described that non-AUG can be used as a start codon for protein translation (Andrews and Rothnagel 2014; Belinky Rogozin and Koonin 2017; Cairns, DeMaria, Poulin, Sancho, Liu, Zhang, Campos-Rivera, Karey and Estes 2011; Gao, Wan, Liu, Ma, Shen and Qian 2015; Ingolia Lareau and Weissman 2011; Ivanov, Firth, Michel, Atkins and Baranov 2011; Lee, Liu, Lee, Huang, Shen and Qian 2012; Ma, Ward, Jungreis, Slavoff, Schwaid, Neveu, Budnik, Kellis and Saghatelian 2014; Mehdi Ono and Gupta 1990; Peabody 1989; Slavoff, Mitchell, Schwaid, Cabili, Ma, Levin, Karger, Budnik, Rinn and Saghatelian 2013; Zhu, Orre, Johansson, Huss, Boekel, Vesterlund, Fernandez-Woodbridge, Branca and Lehtio 2018). Therefore, in predicting the coding potential of RNA, in addition to selecting AUG as the start codon, a triplet can also be used as the start codon. In 2012, Lee *et al.* develop a technology for global translation initiation sequencing (GTI-seq) (Lee, Liu, Lee, Huang, Shen and Qian 2012). The GTI-seq results reveal that almost half of the codon at downstream translation initiation sites (downstream TIS, dTIS) still use AUG as the main initiator. However, half of the total TIS codons are upstream TIS codons (uTISs), and 74.4% of the uTIS codons are non-AUG codons (Gao, Wan, Liu, Ma, Shen and Qian 2015; Lee, Liu, Lee, Huang, Shen and Qian 2012). CUG is the most prominent codon among the cognate codons of AUG (Dever 2012; Kearse and Wilusz 2017), and the frequency of occupation is even higher than that of AUG with 30.3% vs 25.6% (Lee, Liu, Lee, Huang, Shen and Qian 2012). What is the difference between the initiation of non-AUG codons and AUG initiation, and which factors can regulate the choice of initiation site, is still an open question. Although the synthesis of most proteins starts with the classical AUG codon, the initiation at the codons of CUG and GUG is also ubiquitous and biologically significant. The non-AUG initiation significantly affects many aspects of translation (Ingolia Lareau and Weissman 2011; Mackowiak, Zauber, Bielow, Thiel, Kutz, Calviello, Mastrobuoni, Rajewsky, Kempa and Selbach et al. 2015). In 2011, research by Ingolia *et al.* show that many TIS codons, especially uTIS codons, occur at the non-AUG codons (Ingolia Lareau and Weissman 2011). In 2012, Slavoff *et al.* use mass spectrometry in human cells to verify 90 SEPs, of which 86 were new peptides, and most of them use non-AUG as the start codon (Slavoff, Mitchell, Schwaid, Cabili, Ma, Levin, Karger, Budnik, Rinn and Saghatelian 2013). They also point out that if there is no upstream AUG, the initiation codon is assigned to a frame near-cognate codons embedded in the Kozak consensus sequence, and the Kozak sequence plays an important role at the non-AUG start sites (Ingolia Lareau and Weissman 2011; Kozak 1986). Study by Zhu *et al.* find that most of the sequences using non-AUG as the start codon contained a strong Kozak motif, which indicates that the near-cognate codons in the strong Kozak sequence may be a prerequisite for uORF translation (Kozak 1986; Kozak 1987; Zhu, Orre, Johansson, Huss, Boekel, Vesterlund, Fernandez-Woodbridge, Branca and Lehtio 2018). In addition, a strong RNA secondary structure starting approximately 15nt downstream of the start site can significantly improve the initiation efficiency of non-AUG codons (Kozak 1990).

In recent years, databases of sORFs have been successively constructed. In 2015, Olexiouk *et al.* analysis and identify the ribosome profiling data, and build sORFs.org database, which contains sORF data of human, mouse and fruit fly (Olexiouk, Crappe, Verbruggen, Verhegen, Martens and Menschaert 2016). In 2017, they update the database and add the sORF data of zebrafish, rat and C. elegans (Olexiouk Van Criekinge and Menschaert 2018). In continue, a database of SEPs, SmProt (Hao, Zhang, Niu, Cai, Luo, He, Zhang, Zhang, Qin and Yang et al. 2018), is established, which includes 255 010 SEPs derived from 291 cell lines and tissues from eight common species. At the same year, the ARA-PEPs database is constructed by Hazarika *et al.*, which collects 13 748 SEPs in arabidopsis, all of them are verified at the transcription level(Hazarika, De Coninck, Yamamoto, Martin, Cammue and van Noort 2017). The upstream ORF (uORF) is a type of sORF. In eukaryotes, the uORF is located in the 5’-untranslated region (5’UTR) of mRNA, which can regulate the translation of downstream ORFs (dORFs), thereby regulating gene expression (Lovett and Rogers 1996; Meijer and Thomas 2002; Morris and Geballe 2000; Oliveira and McCarthy 1995; Vilela and McCarthy 2003). The uORF is widely presented in eukaryotes (Calvo Pagliarini and Mootha 2009; Crowe Wang and Rothnagel 2006; Hayden and Bosco 2008; Somers Poyry and Willis 2013; von Arnim Jia and Vaughn 2014). Approximately 50% of human genes contain uORF in their 5’UTR, and when uORF is presented, it results in reducing expression of subsequent proteins (Calvo Pagliarini and Mootha 2009). In 2014, Wethmar *et al.* construct the uORF database, uORFdb, which is a database of uORF and manually find from all relevant literature listed in the PubMed (Wethmar, Barbosa-Silva, Andrade-Navarro and Leutz 2014). We used the sORF data from the sORFs.org database (Olexiouk Van Criekinge and Menschaert 2018; Olexiouk, Crappe, Verbruggen, Verhegen, Martens and Menschaert 2016) to analyze the start codons (Figure 1). The results showed that the classical start codon AUG accounted for only 16.79%, while GUG accounted for 40.63%, and CUG and UUG accounted for 22.11% and 19.04%, which indicated that the non-ATG start codon accounts for a large part of sORF translation.

**Figure 1.**
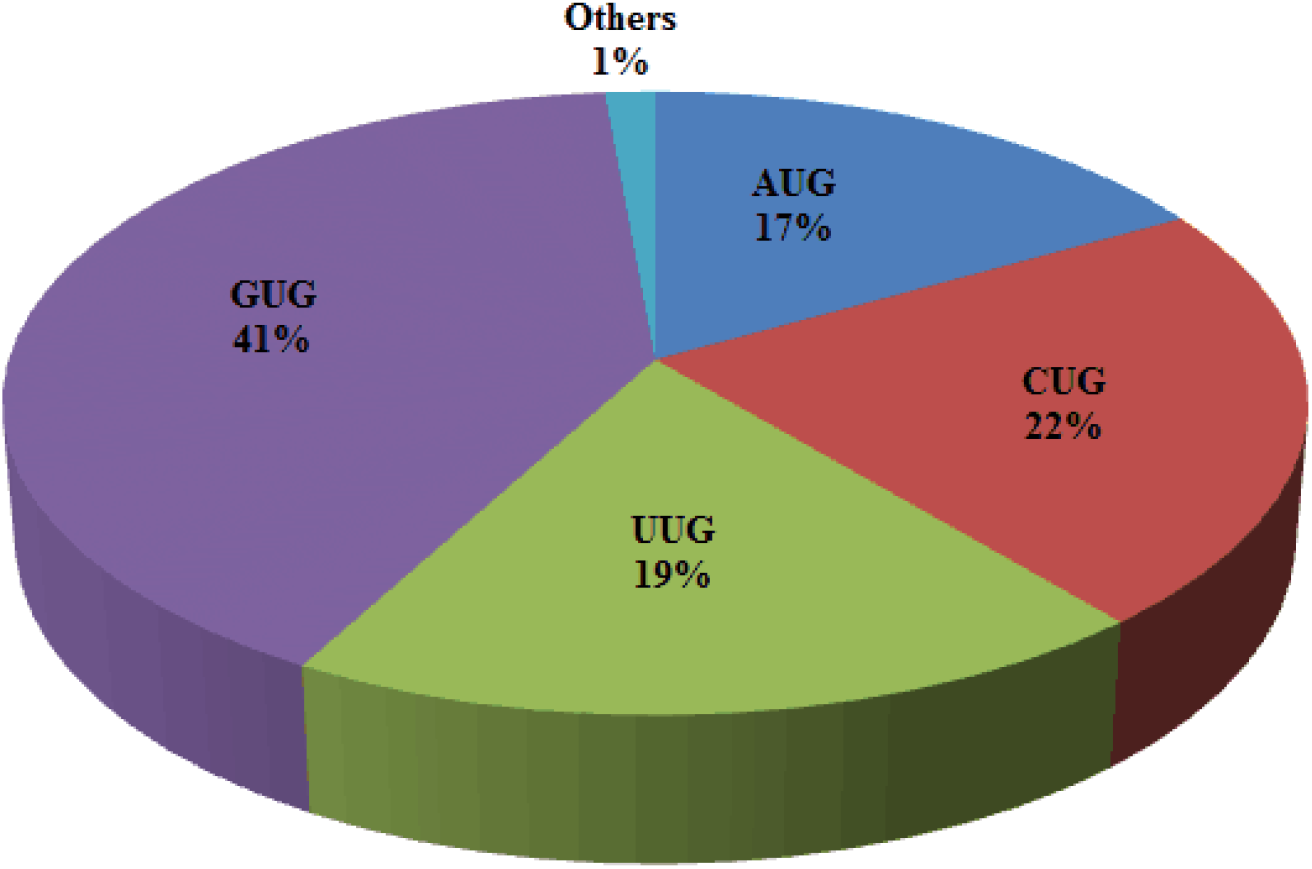
Composition of the start codons on the sORF data from sORFs.org. The data used includes 8512 sORFs, of which AUG with 16.79%, GUG and CUG with 40.63% and 22.11%, UUG with 19.04%

Based on the above analysis, we think that in addition to using the classical AUG codon as the start codon, we can also use near-cognate codons as the start codon to build a model to predict the coding potential of RNA. Here, we consider CUG and GUG as the non-AUG start codons. The two codons are the most widely distributed in research (Belinky Rogozin and Koonin 2017; Ingolia Lareau and Weissman 2011; Lee, Liu, Lee, Huang, Shen and Qian 2012; Ma, Ward, Jungreis, Slavoff, Schwaid, Neveu, Budnik, Kellis and Saghatelian 2014; Slavoff, Mitchell, Schwaid, Cabili, Ma, Levin, Karger, Budnik, Rinn and Saghatelian 2013). In 2012, researchers found that CUG can be used as a start codon to translate proteins (Dever 2012; Starck, Jiang, Pavon-Eternod, Prasad, McCarthy, Pan and Shastri 2012). By including the non-AUG start codons, the search parameters of the sORF can be made less stringent. This is an important factor to consider, because the non-AUC is pretty common as the start codon. The polypeptides encoded using non-AUG as the start codon have been identified by mass spectrometry (MS) (Slavoff, Mitchell, Schwaid, Cabili, Ma, Levin, Karger, Budnik, Rinn and Saghatelian 2013). Most previous algorithms searching for mRNA were limited to more than 300 nucleotides, which translate at least 100 amino acids (Hu, Xu, Hu and Lu 2017; Kang, Yang, Kong, Hou, Meng, Wei and Gao 2017; Li Zhang and Zhou 2014; Sun, Chen, Jiang, Song, Wang and Sun 2013; Wang, Park, Dasari, Wang, Kocher and Li 2013; Wucher, Legeai, Hedan, Rizk, Lagoutte, Leeb, Jagannathan, Cadieu, David and Lohi et al. 2017). This defect trigger the classical gene annotation software to misclassify some ncRNA containing sORF. In 2010, Hanada *et al.* propose a program specially designed to predict sORF, sORF finder, which uses hexamer features to predict the coding potential of sORF, but the prediction results obtained using only one feature will have a high false positive rate (Hanada, Akiyama, Sakurai, Toyoda, Shinozaki and Shiu 2010). In 2018, Hill *et al.* develop a model of mRNN based on Recurrent Neural Networks (RNN), and they predict the coding potential of sORF (Hill, Kuintzle, Teegarden, Merrill, Danaee and Hendrix 2018). In November 2019, Zhu *et al.* also develop a tool specifically for predicting micropeptides, MiPepid, which is a tool developed by using K-mer features and machine learning (Zhu and Gribskov 2019). We tested it by using human sORF data (Tong and Liu 2019) and found that the false positives of MiPepid were high. A large amount of scientific literature has reported that the prediction of sORF is still an open question (Basrai Hieter and Boeke 1997; Hanada, Zhang, Borevitz, Li and Shiu 2007; Wang, Li, Zhang, Zheng, Xu, Ye, Yu and Wong 2003). Moreover, sORF can start translation with non-AUG, which also brings new challenges to the research. Therefore, it is necessary to develop a tool that can accurately predict the coding potential of sORF to accurately distinguish whether RNA containing sORF can encode peptide. Thus, in the study of this article, we used AUG, GUG and CUG as the start codons, and updated our tool CPPred (Tong and Liu 2019) to CPPred-sORF for sORF data.

## METHODS

### Dataset

As of 2 April 2019, we downloaded the coding RNAs and ncRNAs of 7 model organisms of human, mouse, zebrafish, fruit fly, nematode, yeast and arabidopsis from RefSeq. Finally, we obtained 19,707 non-redundant lncRNAs and 25,407 non-redundant small coding RNAs, as shown in Figure 2. Compared with the recognition of both small coding RNA and small ncRNA, lncRNA and small coding RNA are more difficult to distinguish, because the longer the sequence, the easier to contain ORF, that is to say, it is more likely to have the ability to encode protein. The first step of data construction is to download positive and negative data. We downloaded human, mouse, zebrafish, fruit fly, nematode, yeast and arabidopsis ncRNA, coding RNA sequences and their corresponding with CDS sequences from RefSeq, and we obtained 507,418 coding RNAs and 54,947 ncRNAs. The second step is to construct small coding RNAs and lncRNAs. For the construction of small coding RNAs, we use the length of 303nt as the cutoff and use the CDS corresponding to the coding RNA, if the length of CDS is less than or equal to 303nt, then it is saved, and finally get 326,414 small coding RNAs. For the lncRNA, we use 200nt as the threshold. If the sequence length is 200nt or more, which is retained, and we finally obtain 24,837 lncRNAs. The third step is the operation of removing redundant data, which is different from the order of removing redundant sequences constructed by CPPred dataset (Tong and Liu 2019). Here we first removed the entire data and then build the training set and testing set. We use CD-Hit to reduce redundancy with sequence identity of 80% (Lertampaiporn, Thammarongtham, Nukoolkit, Kaewkamnerdpong and Ruengjitchatchawalya 2014; Liu, Fang, Liu, Wang, Chen and Chou 2015; Sun, Liu, Zhang and Meng 2015; Tong and Liu 2019). In the end, we obtained 25,407 non-redundant small coding RNAs and 19,707 non-redundant lncRNAs. The fourth step is to build training set and testing set. We randomly extracted two-thirds from these non-redundant sequences as the training set, and the remaining sequence as the testing set. For the small coding RNAs, 16,938 and 8469 sequences as training and testing sets, respectively. For the lncRNAs, 13,138 sequences are used as the training set, and 6569 is used as the testing set. This construction method can well ensure that the RNA sequences between the training set and the testing set, and the training set and the testing set are all non-redundant.

**Figure 2.**
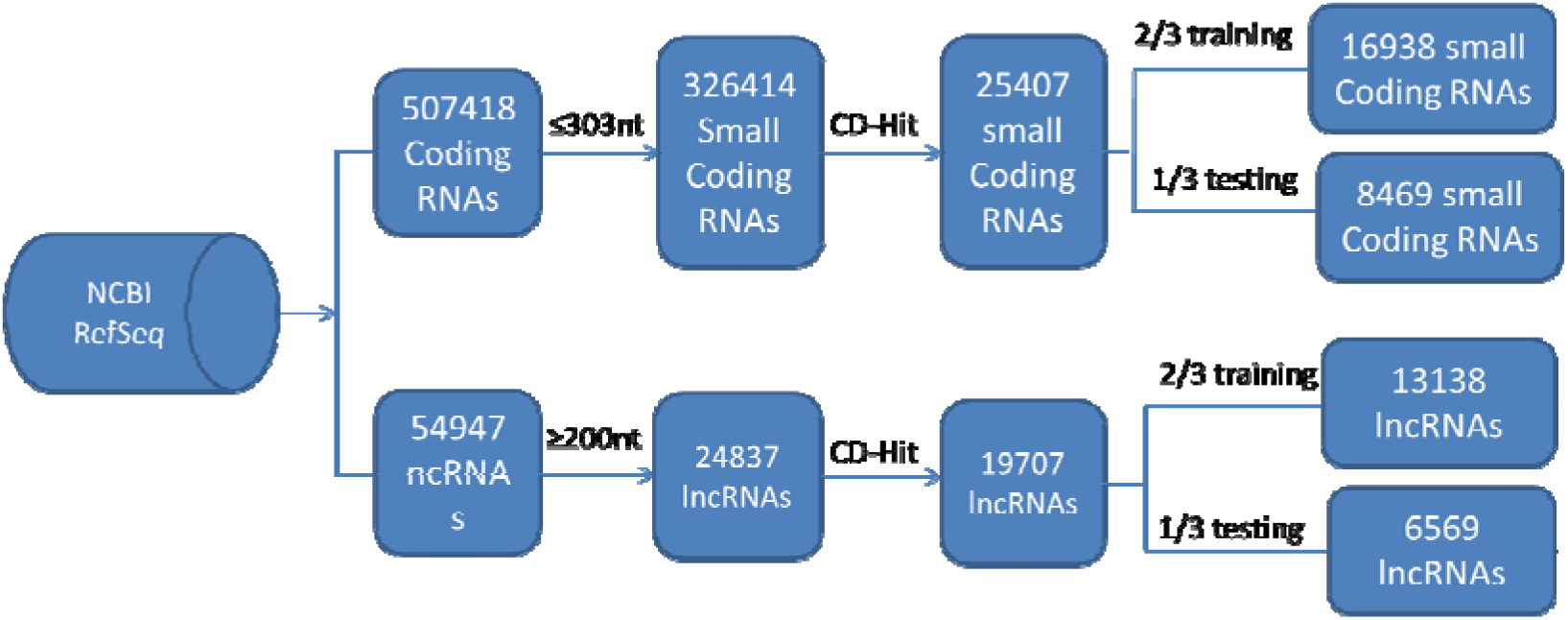
Flow chart of sORF data construction. We downloaded the coding RNAs and ncRNAs of 7 model organisms from the RefSeq. We used the length threshold as a condition, where the thresholds for coding RNA and ncRNA were 303nt and 200nt, respectively, to choose out 326,414 small coding RNAs and 24,837 lncRNAs. The CD-Hit tool was used to remove redundant sequences from small coding RNAs and lncRNAs according to the sequence identity threshold of 80%. Finally, two-thirds were randomly selected as the training set, and the remaining sequences were used as the testing set

In order to test our method, we build testing sets of other species. We downloaded the coding RNAs and ncRNAs of chicken, pig, rabbit, cow, rhesus, maize and soybean from RefSeq. Based on the same construction method, that is, the small coding RNA and lncRNA are extracted based on the sequence length, and then CD-Hit is used to delete redundant sequences according to the sequence identity threshold of 80%, and finally the non-redundant sequence is obtained between the training set constructed above. Because these species are not as extensive as those studied by model organisms, the resulting non-redundant sequences, especially the lncRNAs, are particularly rare. For example, there are only 50 non-redundant lncRNA sequences for cattle, 11 for chickens, 9 for lncRNAs for pigs, 5 for lncRNAs for rhesus monkeys and none for rabbits. Relatively speaking, only the sequences of maize and soybean are relatively large, so here we list the maize and soybean data (Table 1).

**Table 1.** The non-redundant small coding RNA and lncRNA of maize and soybean

| Organism | small coding RNAs | lncRNAs |
| --- | --- | --- |
| maize | 315 | 281 |
| soybean | 315 | 110 |

### CPPred-sORF Features

In the CPPred(Tong and Liu 2019), each RNA sequence is encoded into 38-dimensional features, which includes ORF length, ORF coverage, ORF integrity, Fckett score, Hexamer score, pI, Gravy, instability and CTD features. In CPPred-sORF, we added the GC count (GCcount) and 11 important codon content (mRNN-11codons) features obtained from the work of mRNN. Among them, the GCcount has been extensively studied for the prediction of coding potential (Biswas, Zhang, Wu and Gao 2013; Brocchieri, Kledal, Karlin and Mocarski 2005; Hu, Xu, Hu and Lu 2017; Omasits, Varadarajan, Schmid, Goetze, Melidis, Bourqui, Nikolayeva, Quebatte, Patrignani and Dehio et al. 2017; Pohl Theissen and Schuster 2012; Wang, Yin, Li, Yu, Wang, Xu, Cao, Bao, Wang and Abbasi et al. 2019). In 2018, Hill *et al.* analysis that eleven codons (UAC, AAC, UAU, AUC, UUC, GAG, AAG, GAU, GAC, AAU and GUG) play a very important role in predicting the coding potential of RNA (Hill, Kuintzle, Teegarden, Merrill, Danaee and Hendrix 2018). We use these two features because they are recognizable for distinguishing coding RNA sequences from non-coding RNAs (Figure 3A and Figure 3B). In addition, more importantly, for the use of the start codon, in addition to the AUG as the start codon, CUG and GUG are also used as the start codon (Belinky Rogozin and Koonin 2017; Ingolia Lareau and Weissman 2011; Lee, Liu, Lee, Huang, Shen and Qian 2012; Ma, Ward, Jungreis, Slavoff, Schwaid, Neveu, Budnik, Kellis and Saghatelian 2014; Slavoff, Mitchell, Schwaid, Cabili, Ma, Levin, Karger, Budnik, Rinn and Saghatelian 2013). The start codons used are different, and the six characteristics of ORF length, ORF coverage, ORF integrity, pI, Gravy and instability will change accordingly.

**Figure 3.**
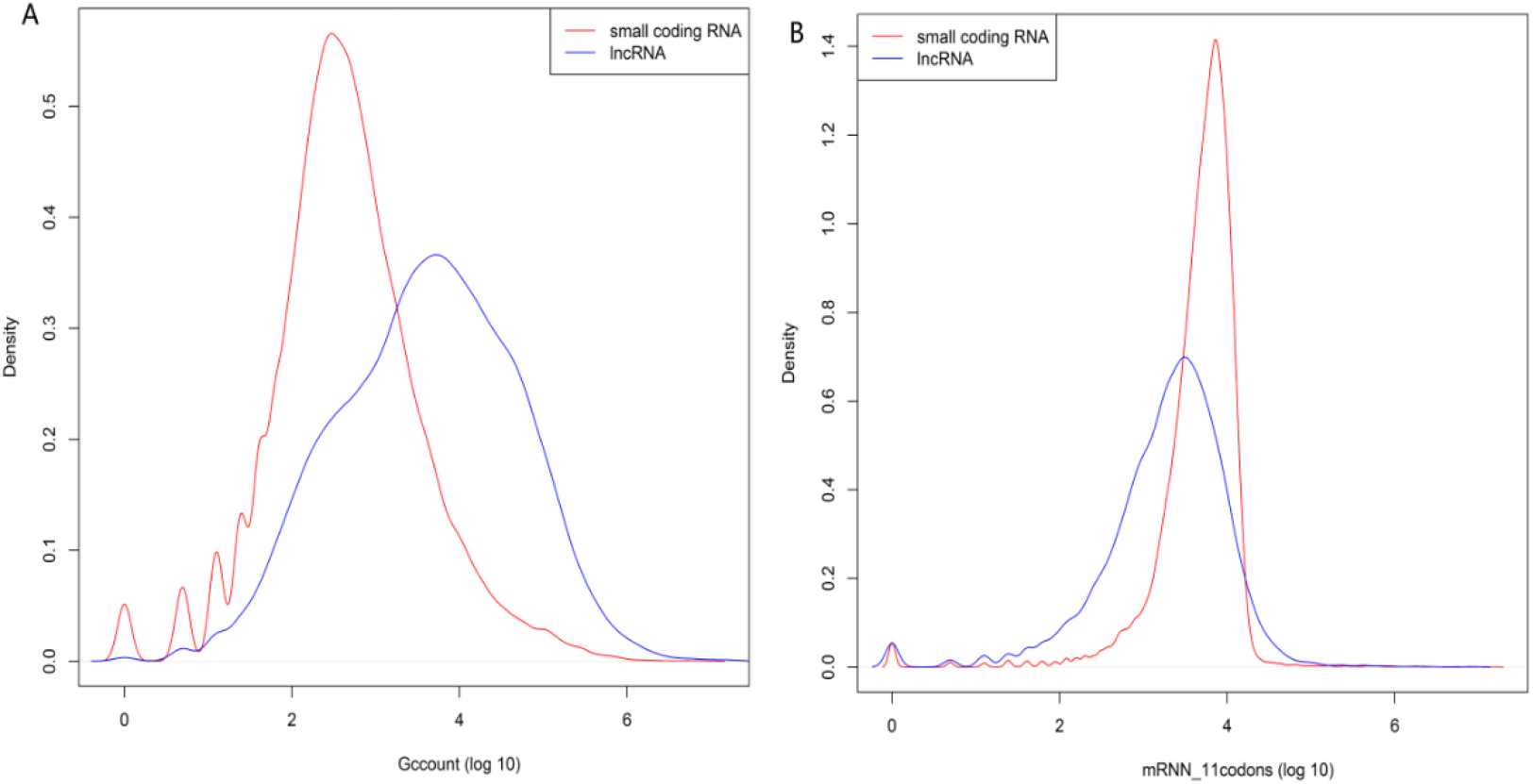
Score distribution between small coding RNAs and lncRNAs for Gccount (A) and mRNN-11codons (B). The training set containing 16938 small coding RNAs and 13138 lncRNAs are used

### Feature selection

The purpose of feature selection is to reduce useless features, make the model more general, and reduce the risk of overfitting. In our previous work CPPred, we use mRMR-IFS method to select the best feature subset (Tong and Liu 2019). In CPPred-sORF, we still use the mRMR-IFS method to select the best feature subset. After obtaining 40 ordered features according to mRMR, and then adding features one by one using IFS, 40 feature subsets can be obtained. For these 40 feature subsets, we obtain corresponding feature sets from the training set. Then, based on the results of 10-fold cross-validation on the training set, the best feature subset is selected and applied to the entire training set to obtain the final model.

### SVM classifier and evaluation method

As with the SVM classifier used by CPPred (Tong and Liu 2019), here we use Libsvm (libsvm-3.22) to classify and predict small coding RNAs and lncRNAs. The radial basis function (RBF) is selected as the kernel function, the MCC value is served as an index of the optimization parameter, and the (C, γ) pair with the largest MCC value in cross-validation is selected. Here, the best C = 7.29377990848 and γ = 3.32495158456 can be obtained by using the top 39 features. Same as CPPred (Tong and Liu 2019), we use SN, SP, ACC, PRE, F-score, AUC and MCC to evaluate the performance of CPPred-sORF.

## RESULTS AND DISCUSSION

In this article, we developed the CPPred-sORF, a prediction tool for the coding potential of sORF. In addition to the ORF length, ORF coverage, ORF integrity, Fickett score, Hexamer score, pI, Gravy, instability and CTD features. In CPPred-sORF, we added the features of GCcount and mRNN-11codons. Especially, for the start codon, it includes the start codons CUG and GUG in addition to AUG. Moreover, we use small coding RNAs and lncRNAs as research objects to predict the coding potential.

### Feature selection results using mRMR-IFS method

Removing some useless or redundant features can not only improve the prediction ability of the model, but also avoid overfitting. The mRMR-IFS is used for feature selection. First, the mRMR was applied to sort the 40-dimensional features of the training set, and then added the sorted features one by one with the IFS. We use 10-fold cross-validation for training and testing to draw feature selection curve. As shown in Figure 4, the best prediction performance is obtained when using the top sorted 39 features, with an MCC value of 0.770. Therefore, in order to predict the coding potential of RNA, we selected the top 39 features as the best feature set to construct the final model.

**Figure 4.**
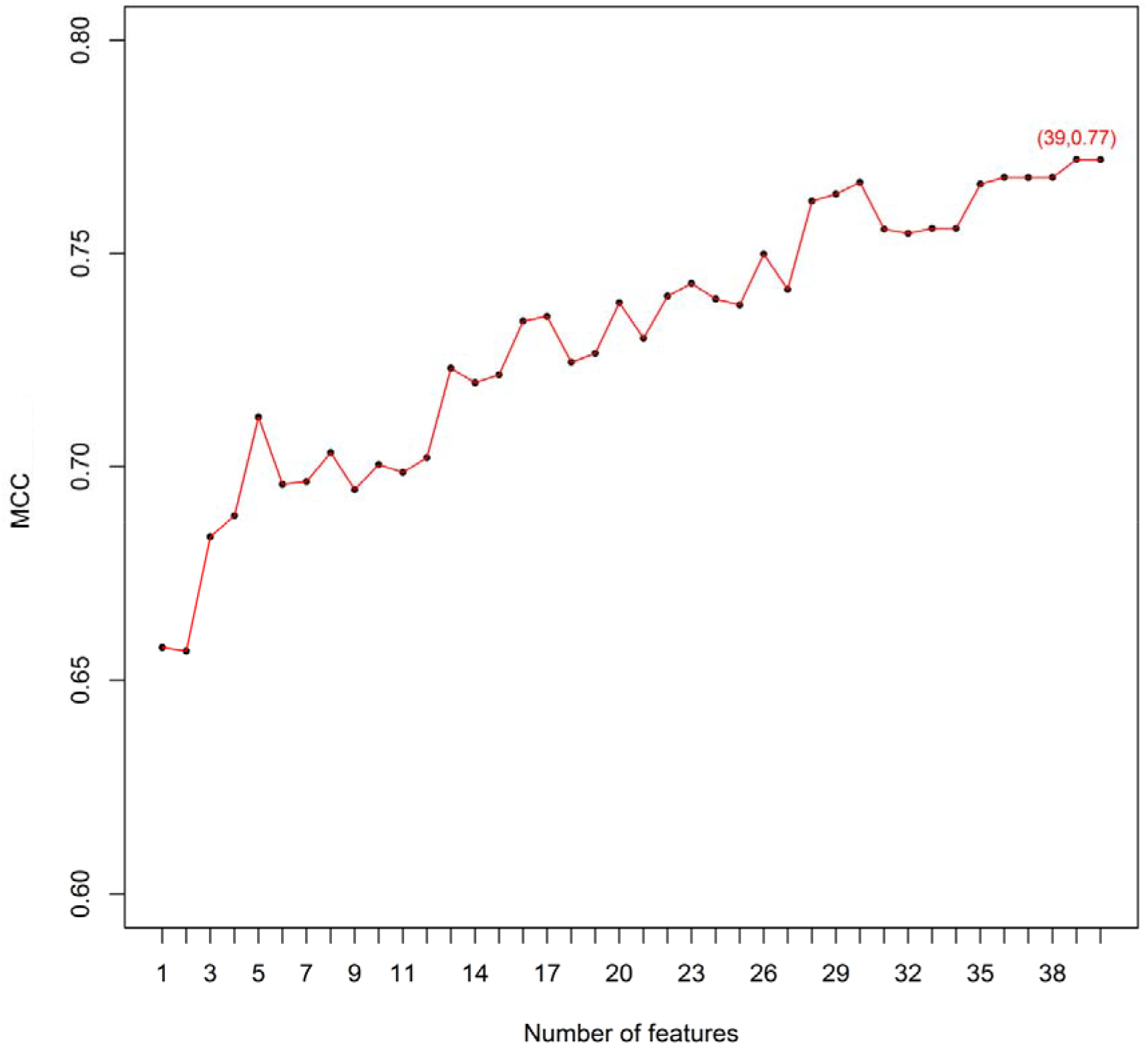
Feature selection curve. The number of features is the horizontal coordinate, and the vertical coordinate is the MCC of corresponding 10-fold cross-validated

In addition, we see that when the features are added one by one, the MCC value does not increase steadily, but fluctuates up and down. After adding the second, sixth, ninth, eleventh, fourteenth, eighteenth, twenty-first, twenty-fourth, twenty-fifth, twenty-seventh, thirty-first, thirty-fourth feature, the MCC value not only did not increase, but decreased, and we will also discuss this point below. Moreover, we also found that when only the first feature is used, that is, only the ORF coverage is used (Table 2), the MCC reaches 0.66, which is only 0.11 different from the maximum MCC value of 0.77. This shows that ORF coverage plays a very important role in distinguishing small coding RNAs from lncRNAs. The ORF length related to ORF coverage ranks fourth in the ranking of mRMR, which is also ranked high. This shows that the features of ORF-related attributes critical role in predicting the coding potential of RNA.

**Table 2.** Features ranked using mRMR

| Order | Feature | Score | Order | Feature | Score | Order | Feature | Score |
| --- | --- | --- | --- | --- | --- | --- | --- | --- |
| 1 | Coverage | 0.411 | 15 | G4 | 0.001 | 29 | G0 | -0.007 |
| 2 | pI | 0.007 | 16 | mRNN-11codons | 0.003 | 30 | Gravy | -0.007 |
| 3 | Hexamer | 0.035 | 17 | AC | 0.001 | 31 | AT | -0.008 |
| 4 | Length | 0.013 | 18 | T3 | -0.000 | 32 | A3 | -0.009 |
| 5 | Fickett | 0.040 | 19 | C3 | -0.001 | 33 | T2 | -0.010 |
| 6 | C0 | 0.011 | 20 | A0 | -0.001 | 34 | C | -0.014 |
| 7 | C4 | 0.007 | 21 | Instability | -0.001 | 35 | C2 | -0.015 |
| 8 | T1 | 0.005 | 22 | A1 | -0.001 | 36 | G2 | -0.017 |
| 9 | TG | 0.005 | 23 | Integrity | -0.002 | 37 | A | -0.019 |
| 10 | GCcount | 0.006 | 24 | G1 | -0.002 | 38 | A2 | -0.019 |
| 11 | T4 | 0.003 | 25 | AG | -0.002 | 39 | G | -0.021 |
| 12 | TC | 0.004 | 26 | T0 | -0.004 | 40 | T | -0.026 |
| 13 | GC | 0.002 | 27 | C1 | -0.005 |  |  |  |
| 14 | A4 | 0.001 | 28 | G3 | -0.007 |  |  |  |

### CPPred-sORF results on independent testing set

Based on the feature selection results of the 10-fold cross-validation on the training set, we finally select the top 39 features to apply to the entire training set, and then input them into the SVM classifier to obtain the final prediction model (39Features). Finally, the model is tested and evaluated on independent tests. We compared CPPred-sORF with CPPred (Tong and Liu 2019), CPAT (Wang, Park, Dasari, Wang, Kocher and Li 2013), CPC2 (Kang, Yang, Kong, Hou, Meng, Wei and Gao 2017), PLEK (Li Zhang and Zhou 2014), sORF finder (Hanada, Akiyama, Sakurai, Toyoda, Shinozaki and Shiu 2010) and mRNN (Hill, Kuintzle, Teegarden, Merrill, Danaee and Hendrix 2018). Among these tools, sORF finder was specially developed for the prediction of sORF. When the mRNN was developed, the situation of sORF was also considered. Therefore, The CPPred-sORF and these two softwares will be more convincing.

From Table 3, we can see that CPPred-sORF outperform CPPred, CPAT, CPC2, PLEK, mRNN and sORF finder on the independent testing set. Their ACC values are 88.49% versus 85.84%, 54.51%, 45.44% 41.46%, 73.84% and 54.85%, and their MCC values are 0.768, 0.730, 0.228, 0.034, −0.112, 0.492 and 0.054. Overall, CPPred-sORF has an obvious advantage in predicting small coding RNAs and lncRNAs. Among them, the CPAT, CPC2, and PLEK are not specifically trained and tested on small coding RNAs, so it is understandable that they have disadvantages in small coding RNAs and lncRNAs prediction.

**Table 3.**
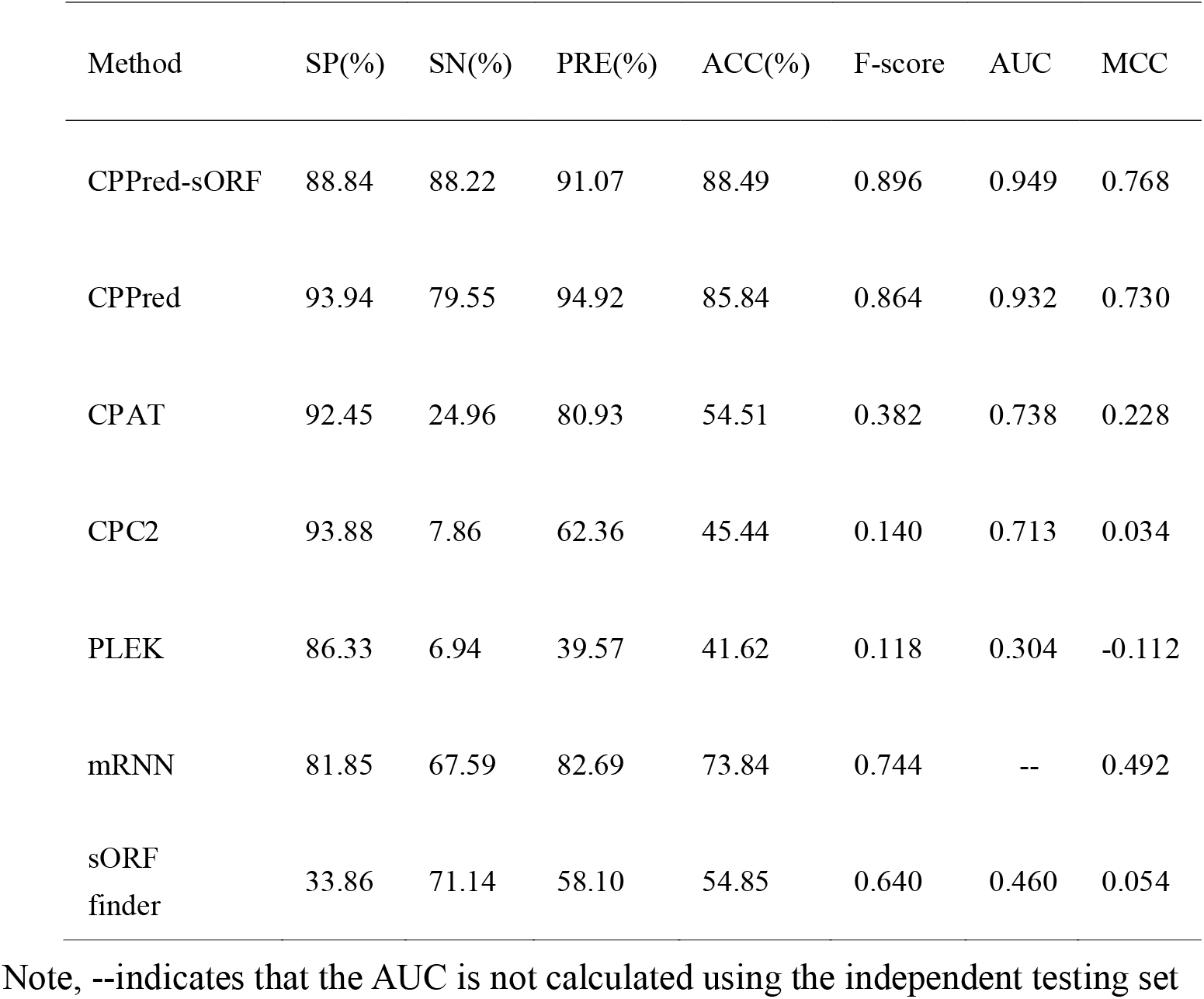
Performance of CPPred-sORF (39Features) and CPPred (Integrated-Model), CPAT, CPC2, PLEK, mRNN and sORF finder on independent testing set

| Method | SP(%) | SN(%) | PRE(%) | ACC(%) | F-score | AUC | MCC |
| --- | --- | --- | --- | --- | --- | --- | --- |
| CPPred-sORF | 88.84 | 88.22 | 91.07 | 88.49 | 0.896 | 0.949 | 0.768 |
| CPPred | 93.94 | 79.55 | 94.92 | 85.84 | 0.864 | 0.932 | 0.730 |
| CPAT | 92.45 | 24.96 | 80.93 | 54.51 | 0.382 | 0.738 | 0.228 |
| CPC2 | 93.88 | 7.86 | 62.36 | 45.44 | 0.140 | 0.713 | 0.034 |
| PLEK | 86.33 | 6.94 | 39.57 | 41.62 | 0.118 | 0.304 | -0.112 |
| mRNN | 81.85 | 67.59 | 82.69 | 73.84 | 0.744 | -- | 0.492 |
| sORF<br>finder | 33.86 | 71.14 | 58.10 | 54.85 | 0.640 | 0.460 | 0.054 |

Note, --indicates that the AUC is not calculated using the independent testing set

We can see from Table 4 and Table 5 that the CPPred-sORF is still superior to other prediction tools on maize and soybean testing sets. The performance of CPPred-sORF on the maize testing set is better than that of the soybean testing set. This may be because the positive and negative data in the maize testing set is almost balanced, while the positive data in the soybean testing set is almost three times the negative data. The unbalanced data may degrade the test capability (Zheng, Zhang, Zhao, Tong, Hong, Xie and Liu 2018).

**Table 4.** Performance of CPPred-sORF, CPPred (Integrated-Model), CPAT, CPC2, PLEK, mRNN and sORF finder on maize testing set

| Method | SP(%) | SN(%) | PRE(%) | ACC(%) | F-score | AUC | MCC |
| --- | --- | --- | --- | --- | --- | --- | --- |
| CPPred-sORF | 80.78 | 89.52 | 83.93 | 85.40 | 0.866 | 0.925 | 0.708 |
| CPPred | 83.63 | 67.30 | 82.17 | 75.00 | 0.740 | 0.844 | 0.513 |
| CPAT | 68.33 | 30.25 | 51.63 | 48.24 | 0.382 | 0.599 | -0.015 |
| CPC2 | 86.48 | 10.79 | 47.22 | 46.48 | 0.176 | 0.646 | -0.042 |
| PLEK | 77.58 | 7.30 | 26.74 | 40.44 | 0.115 | 0.387 | -0.215 |
| mRNN | 48.04 | 55.73 | 54.52 | 52.10 | 0.551 | -- | 0.038 |
| sORF<br>finder | 24.20 | 95.56 | 58.56 | 61.91 | 0.726 | 0.532 | 0.286 |

**Table 5.** Performance of CPPred-sORF, CPPred (Integrated-Model), CPAT, CPC2, PLEK, mRNN and sORF finder on soybean testing set

| Method | SP(%) | SN(%) | PRE(%) | ACC(%) | F-score | AUC | MCC |
| --- | --- | --- | --- | --- | --- | --- | --- |
| CPPred-sORF | 76.36 | 89.52 | 91.56 | 86.12 | 0.905 | 0.875 | 0.646 |
| CPPred | 83.64 | 67.30 | 92.17 | 71.53 | 0.778 | 0.853 | 0.448 |
| CPAT | 92.73 | 30.25 | 92.23 | 46.46 | 0.456 | 0.859 | 0.235 |
| CPC2 | 90.00 | 10.79 | 75.56 | 31.29 | 0.189 | 0.837 | 0.011 |
| PLEK | 94.55 | 7.30 | 79.31 | 29.88 | 0.134 | 0.756 | 0.032 |
| mRNN | 83.64 | 55.73 | 90.67 | 62.97 | 0.690 | -- | 0.347 |
| sORF<br>finder | 59.09 | 95.56 | 86.99 | 86.12 | 0.911 | 0.770 | 0.615 |

On the maize testing set, the MCC value of CPPred-sORF is 0.708, which is higher than other prediction tools. The CPPred and mRNN have similar MCC values, which are better than CPAT, CPC2, and PLEK. This may be because the training sets of CPPred and mRNN both include small coding RNAs to build prediction models. However, The CPAT, CPC2, and PLEK are aimed at predicting the coding potential of longer RNAs, so they have disadvantages in predicting small coding RNAs, which may also be the reason why their performance is relatively different from other tools. We can also find that the MCC value of the sORF finder tool is worse than CPPred-sORF, but it is superior to other tools. This is because the sORF finder is a prediction tool specifically for sORF. In addition, as shown in Table 3, Table 4 and Table 5, we can see that although the SN of sORF finder is higher than other prediction tools, its SP is much worse than other prediction tools. This indicates that many lncRNAs are predicted to coding RNA, and the false positives are higher than other tools. This may be because the sORF finder is constructed using only one feature, so its false positives are high.

### Analysis of CPPred-sORF Features

From the selection feature curve in Figure 4, we can see that when the sorted features are added one by one, the MCC fluctuates up and down. So we try to remove those features that fluctuate downwards to train a model. We remove the features that MCC drops by 0.01 or more, namely C0, TG, T3, instability, C1, AT and C. Then, compared with the 39 features that obtained the optimal MCC value in Figure 4, we get 32 features. We apply these 32 features to the entire training set and input them to the SVM classifier to obtain a new prediction model (32Features). The test results on the independent testing set, maize testing set and soybean testing set are shown in Table 6.

**Table 6.**
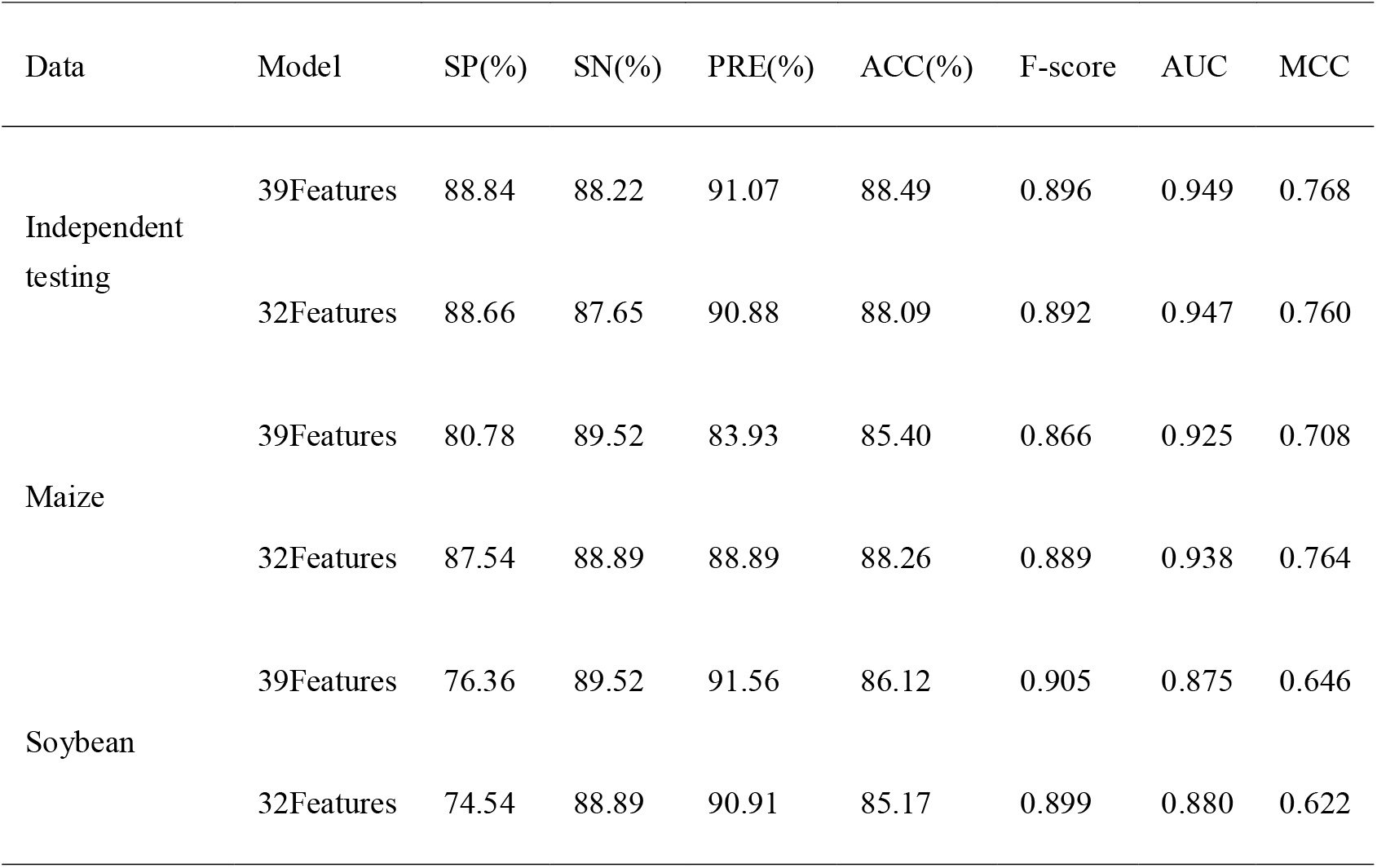
Performance of 39Features and 32Features on independent testing set, maize testing set and soybean testing set

| Data | Model | SP(%) | SN(%) | PRE(%) | ACC(%) | F-score | AUC | MCC |
| --- | --- | --- | --- | --- | --- | --- | --- | --- |
| Independent testing | 39Features | 88.84 | 88.22 | 91.07 | 88.49 | 0.896 | 0.949 | 0.768 |
|  | 32Features | 88.66 | 87.65 | 90.88 | 88.09 | 0.892 | 0.947 | 0.760 |
| Maize | 39Features | 80.78 | 89.52 | 83.93 | 85.40 | 0.866 | 0.925 | 0.708 |
|  | 32Features | 87.54 | 88.89 | 88.89 | 88.26 | 0.889 | 0.938 | 0.764 |
| Soybean | 39Features | 76.36 | 89.52 | 91.56 | 86.12 | 0.905 | 0.875 | 0.646 |
|  | 32Features | 74.54 | 88.89 | 90.91 | 85.17 | 0.899 | 0.880 | 0.622 |

From Table 6, we observe that compared with 39Features, the results on the independent testing set and the soybean testing set show that the performance of 32Features is slightly reduced, which shows that these features still play a role in predicting the coding potential. On the soybean testing set, the performance obtained by the 32Features model has been improved. This example shows that C0, TG, T3, instability, C1, AT, and C are useless features. Based on the results of the above testing sets, we still use the 39Features as the final model. Although the performance is slightly worse to the 32Features on the soybean testing set, the 39Features is the best on the independent testing set and the maize testing set.

### Analysis of Two New Features of CPPed-sORF

In order to analyze the two newly added features, namely the GCcount and the mRNN-11codons. We remove these two features from the 39 features that obtained the best MCC in Figure 4. Then we will get 37 features. We apply these 37 features to the entire training set to obtain another new prediction model (37Features). The test results on the independent testing set, maize testing set and soybean testing set are shown in Table 7.

**Table 7.** Performance of 39Features, 32Features and 37Features on independent testing set, maize testing set and soybean testing set

| Data | Model | SP(%) | SN(%) | PRE(%) | ACC(%) | F-score | AUC | MCC |
| --- | --- | --- | --- | --- | --- | --- | --- | --- |
| Independent testing | 39Features | 88.84 | 88.22 | 91.07 | 88.49 | 0.896 | 0.949 | 0.768 |
|  | 32Features | 88.66 | 87.65 | 90.88 | 88.09 | 0.892 | 0.947 | 0.760 |
|  | 37Features | 88.86 | 88.00 | 91.06 | 88.38 | 0.895 | 0.947 | 0.765 |
| Maize | 39Features | 80.78 | 89.52 | 83.93 | 85.40 | 0.866 | 0.925 | 0.708 |
|  | 32Features | 87.54 | 88.89 | 88.89 | 88.26 | 0.889 | 0.938 | 0.764 |
|  | 37Features | 80.07 | 88.57 | 83.28 | 84.56 | 0.858 | 0.918 | 0.691 |
| Soybean | 39Features | 76.36 | 89.52 | 91.56 | 86.12 | 0.905 | 0.875 | 0.646 |
|  | 32Features | 74.54 | 88.89 | 90.91 | 85.17 | 0.899 | 0.880 | 0.622 |
|  | 37Features | 75.45 | 88.57 | 91.18 | 85.18 | 0.899 | 0.874 | 0.625 |

As shown in the results of Table 7, we find that the performance of the 39Features is better than the 37Features on the independent testing set, the maize and soybean testing sets, which means that the GCcount and mRNN-11codons can slightly improve the predictive performance of coding potential. This shows that the two new features we added are still useful to distinguish small coding RNAs from lncRNAs.

## CONCLUSION

In this paper, we added the GCcount and mRNN-11codons, and also used CUG and GUG in addition to the classical start codon AUG as the start codon. Our analysis found that these two new features can improve the performance of coding potential prediction. We use 39Features as the final prediction model. On the independent testing set, the MCC and ACC values of CPPred-sORF are significantly higher than CPPred, CPAT, CPC2, PLEK, mRNN, and sORF finder. Their MCCs are 0.768, 0.730, 0.228, 0.034, −0.112, 0.492, and 0.054, respectively and the ACCs are 88.49%, 85.84%, 54.51%, 45.44%, 41.62%, 73.84% and 54.85%. Among them, the MCC and ACC of CPPred and mRNN are better than CPAT, CPC2, PLEK and sORF finder. This may be that the CPPred and mRNN are trained using small coding RNAs when building the models. However, the CPAT, CPC2 and PLEK are targeted to predict long coding RNAs, so they are weaker for predicting small coding RNAs. On the maize and soybean testing sets, the CPPred-sORF is still better than CPPred, CPAT, CPC2, PELK, mRNN and sORF finder. The MCCs on the maize testing set are 0.708, 0.513, −0.015, −0.042, −0.215, 0.038 and 0.286, and the MCCs on the soybean testing set are 0.646 to 0.448, 0.235, 0.011, 0.032, 0.347 and 0.615. The performance of CPPred-sORF on the maize testing set is better than the soybean testing set, which indicates that the balanced and unbalanced data sets may affect prediction performance (Zheng, Zhang, Zhao, Tong, Hong, Xie and Liu 2018). In general, CPPred-sORF has a great advantage in predicting sORF.

## Data availability

Source code was implemented in Python and is freely available at http://www.rnabinding.com/CPPred-sORF.

## Acknowledgements and Funding

We thank the National Supercomputer Center in Guangzhou for the support of computing resources. National Natural Science Foundation of China [31100522]; National High Technology Research and Development Program of China [2012AA020402]; the Fundamental Research Funds for the Central Universities [2016YXMS017]; Special Program for Applied Research on Super Computation of the NSFC-Guangdong Joint Fund (the second phase) [U1501501]. Funding for open access charge: Fundamental Research Funds for the Central Universities [2016YXMS017].

## References

Anderson, D.M., C.A. Makarewich, K.M. Anderson, J.M. Shelton, S. Bezprozvannaya, R. Bassel-Duby, and E.N. Olson. 2016. Widespread control of calcium signaling by a family of SERCA-inhibiting micropeptides. Sci Signal 9: a119.

Anderson, D.M., K.M. Anderson, C.L. Chang, C.A. Makarewich, B.R. Nelson, J.R. McAnally, P. Kasaragod, J.M. Shelton, J. Liou, and R. Bassel-Duby et al. 2015. A micropeptide encoded by a putative long noncoding RNA regulates muscle performance. Cell 160:595–606.

Andrews, S.J. and J.A. Rothnagel. 2014. Emerging evidence for functional peptides encoded by short open reading frames. Nat Rev Genet 15:193–204.

Aspden, J.L., Y.C. Eyre-Walker, R.J. Phillips, U. Amin, M.A. Mumtaz, M. Brocard, and J.P. Couso. 2014. Extensive translation of small Open Reading Frames revealed by Poly-Ribo-Seq. Elife 3:e3528.

Basrai, M.A. P. Hieter and J.D. Boeke. 1997. Small open reading frames: beautiful needles in the haystack. Genome Res 7:768–771.

Bazzini, A.A., T.G. Johnstone, R. Christiano, S.D. Mackowiak, B. Obermayer, E.S. Fleming, C.E. Vejnar, M.T. Lee, N. Rajewsky, and T.C. Walther et al. 2014. Identification of small ORFs in vertebrates using ribosome footprinting and evolutionary conservation. EMBO J 33:981–993.

Belinky, F. I.B. Rogozin and E.V. Koonin. 2017. Selection on start codons in prokaryotes and potential compensatory nucleotide substitutions. Sci Rep 7:12422.

Biswas, A.K., B. Zhang, X. Wu, and J.X. Gao. 2013. CNCTDiscriminator: coding and noncoding transcript discriminator - an excursion through hypothesis learning and ensemble learning approaches. J Bioinform Comput Biol 11:1342002.

Brocchieri, L., T.N. Kledal, S. Karlin, and E.S. Mocarski. 2005. Predicting coding potential from genome sequence: application to betaherpesviruses infecting rats and mice. J Virol 79:7570–7596.

Cairns, V.R., C.T. DeMaria, F. Poulin, J. Sancho, P. Liu, J. Zhang, J. Campos-Rivera, K.P. Karey, and S. Estes. 2011. Utilization of non-AUG initiation codons in a flow cytometric method for efficient selection of recombinant cell lines. Biotechnol Bioeng 108:2611–2622.

Calvo, S.E. D.J. Pagliarini and V.K. Mootha. 2009. Upstream open reading frames cause widespread reduction of protein expression and are polymorphic among humans. Proc Natl Acad Sci U S A 106:7507–7512.

Casson, S.A., P.M. Chilley, J.F. Topping, I.M. Evans, M.A. Souter, and K. Lindsey. 2002. The POLARIS gene of Arabidopsis encodes a predicted peptide required for correct root growth and leaf vascular patterning. Plant Cell 14:1705–1721.

Crowe, M.L. X.Q. Wang and J.A. Rothnagel. 2006. Evidence for conservation and selection of upstream open reading frames suggests probable encoding of bioactive peptides. BMC Genomics 7:16.

Dever, T.E. 2012. Molecular biology. A new start for protein synthesis. Science 336:1645–1646.

D’Lima, N.G., J. Ma, L. Winkler, Q. Chu, K.H. Loh, E.O. Corpuz, B.A. Budnik, J. Lykke-Andersen, A. Saghatelian, and S.A. Slavoff. 2017. A human microprotein that interacts with the mRNA decapping complex. Nat Chem Biol 13:174–180.

Gao, X., J. Wan, B. Liu, M. Ma, B. Shen, and S.B. Qian. 2015. Quantitative profiling of initiating ribosomes in vivo. Nat Methods 12:147–153.

Hanada, K., K. Akiyama, T. Sakurai, T. Toyoda, K. Shinozaki, and S.H. Shiu. 2010. sORF finder: a program package to identify small open reading frames with high coding potential. Bioinformatics 26: 399–400.

Hanada, K., M. Higuchi-Takeuchi, M. Okamoto, T. Yoshizumi, M. Shimizu, K. Nakaminami, R. Nishi, C. Ohashi, K. Iida, and M. Tanaka et al. 2013. Small open reading frames associated with morphogenesis are hidden in plant genomes. Proc Natl Acad Sci U S A 110:2395–2400.

Hanada, K., X. Zhang, J.O. Borevitz, W.H. Li, and S.H. Shiu. 2007. A large number of novel coding small open reading frames in the intergenic regions of the Arabidopsis thaliana genome are transcribed and/or under purifying selection. Genome Res 17:632–640.

Hanyu-Nakamura, K., H. Sonobe-Nojima, A. Tanigawa, P. Lasko, and A. Nakamura. 2008. Drosophila Pgc protein inhibits P-TEFb recruitment to chromatin in primordial germ cells. Nature 451:730–733.

Hao, Y., L. Zhang, Y. Niu, T. Cai, J. Luo, S. He, B. Zhang, D. Zhang, Y. Qin, and F. Yang et al. 2018. SmProt: a database of small proteins encoded by annotated coding and non-coding RNA loci. Brief Bioinform 19:636–643.

Hayden, C.A. and G. Bosco. 2008. Comparative genomic analysis of novel conserved peptide upstream open reading frames in Drosophila melanogaster and other dipteran species. BMC Genomics 9:61.

Hazarika, R.R., B. De Coninck, L.R. Yamamoto, L.R. Martin, B.P. Cammue, and V. van Noort. 2017. ARA-PEPs: a repository of putative sORF-encoded peptides in Arabidopsis thaliana. BMC Bioinformatics 18:37.

Hecht, A., J. Glasgow, P.R. Jaschke, L.A. Bawazer, M.S. Munson, J.R. Cochran, D. Endy, and M. Salit. 2017. Measurements of translation initiation from all 64 codons in E. coli. Nucleic Acids Res 45:3615–3626.

Hill, S.T., R. Kuintzle, A. Teegarden, E.R. Merrill, P. Danaee, and D.A. Hendrix. 2018. A deep recurrent neural network discovers complex biological rules to decipher RNA protein-coding potential. Nucleic Acids Res 46:8105–8113.

Hu, L., Z. Xu, B. Hu, and Z.J. Lu. 2017. COME: a robust coding potential calculation tool for lncRNA identification and characterization based on multiple features. Nucleic Acids Res 45:e2.

Ingolia, N.T. L.F. Lareau and J.S. Weissman. 2011. Ribosome profiling of mouse embryonic stem cells reveals the complexity and dynamics of mammalian proteomes. Cell 147:789–802.

Ivanov, I.P., A.E. Firth, A.M. Michel, J.F. Atkins, and P.V. Baranov. 2011. Identification of evolutionarily conserved non-AUG-initiated N-terminal extensions in human coding sequences. Nucleic Acids Res 39:4220–4234.

Kang, Y.J., D.C. Yang, L. Kong, M. Hou, Y.Q. Meng, L. Wei, and G. Gao. 2017. CPC2: a fast and accurate coding potential calculator based on sequence intrinsic features. Nucleic Acids Res 45:W12–W16.

Kearse, M.G. and J.E. Wilusz. 2017. Non-AUG translation: a new start for protein synthesis in eukaryotes. Genes Dev 31:1717–1731.

Kondo, T., S. Plaza, J. Zanet, E. Benrabah, P. Valenti, Y. Hashimoto, S. Kobayashi, F. Payre, and Y. Kageyama. 2010. Small peptides switch the transcriptional activity of Shavenbaby during Drosophila embryogenesis. Science 329:336–339.

Kozak, M. 1986. Point mutations define a sequence flanking the AUG initiator codon that modulates translation by eukaryotic ribosomes. Cell 44:283–292.

Kozak, M. 1987. An analysis of 5’-noncoding sequences from 699 vertebrate messenger RNAs. Nucleic Acids Res 15:8125–8148.

Kozak, M. 1990. Downstream secondary structure facilitates recognition of initiator codons by eukaryotic ribosomes. Proc Natl Acad Sci U S A 87:8301–8305.

Lauressergues, D., J.M. Couzigou, H.S. Clemente, Y. Martinez, C. Dunand, G. Becard, and J.P. Combier. 2015. Primary transcripts of microRNAs encode regulatory peptides. Nature 520:90–93.

Lee, C., J. Zeng, B.G. Drew, T. Sallam, A. Martin-Montalvo, J. Wan, S.J. Kim, H. Mehta, A.L. Hevener, and R. de Cabo et al. 2015. The mitochondrial-derived peptide MOTS-c promotes metabolic homeostasis and reduces obesity and insulin resistance. Cell Metab 21:443–454.

Lee, S., B. Liu, S. Lee, S.X. Huang, B. Shen, and S.B. Qian. 2012. Global mapping of translation initiation sites in mammalian cells at single-nucleotide resolution. Proc Natl Acad Sci U S A 109:E2424–E2432.

Lertampaiporn, S., C. Thammarongtham, C. Nukoolkit, B. Kaewkamnerdpong, and M. Ruengjitchatchawalya. 2014. Identification of non-coding RNAs with a new composite feature in the Hybrid Random Forest Ensemble algorithm. Nucleic Acids Res 42:e93.

Li, A. J. Zhang and Z. Zhou. 2014. PLEK: a tool for predicting long non-coding RNAs and messenger RNAs based on an improved k-mer scheme. BMC Bioinformatics 15:311.

Liu, B., L. Fang, F. Liu, X. Wang, J. Chen, and K.C. Chou. 2015. Identification of real microRNA precursors with a pseudo structure status composition approach. PLoS One 10:e121501.

Lovett, P.S. and E.J. Rogers. 1996. Ribosome regulation by the nascent peptide. Microbiol Rev 60: 366–385.

Ma, J., C.C. Ward, I. Jungreis, S.A. Slavoff, A.G. Schwaid, J. Neveu, B.A. Budnik, M. Kellis, and A. Saghatelian. 2014. Discovery of human sORF-encoded polypeptides (SEPs) in cell lines and tissue. J Proteome Res 13:1757–1765.

Mackowiak, S.D., H. Zauber, C. Bielow, D. Thiel, K. Kutz, L. Calviello, G. Mastrobuoni, N. Rajewsky, S. Kempa, and M. Selbach et al. 2015. Extensive identification and analysis of conserved small ORFs in animals. Genome Biol 16:179.

Magny, E.G., J.I. Pueyo, F.M. Pearl, M.A. Cespedes, J.E. Niven, S.A. Bishop, and J.P. Couso. 2013. Conserved regulation of cardiac calcium uptake by peptides encoded in small open reading frames. Science 341:1116–1120.

Matsumoto, A., A. Pasut, M. Matsumoto, R. Yamashita, J. Fung, E. Monteleone, A. Saghatelian, K.I. Nakayama, J.G. Clohessy, and P.P. Pandolfi. 2017. mTORC1 and muscle regeneration are regulated by the LINC00961-encoded SPAR polypeptide. Nature 541:228–232.

Mehdi, H. E. Ono and K.C. Gupta. 1990. Initiation of translation at CUG, GUG, and ACG codons in mammalian cells. Gene 91:173–178.

Meijer, H.A. and A.A. Thomas. 2002. Control of eukaryotic protein synthesis by upstream open reading frames in the 5’-untranslated region of an mRNA. Biochem J 367:1–11.

Morris, D.R. and A.P. Geballe. 2000. Upstream open reading frames as regulators of mRNA translation. Mol Cell Biol 20:8635–8642.

Nelson, B.R., C.A. Makarewich, D.M. Anderson, B.R. Winders, C.D. Troupes, F. Wu, A.L. Reese, J.R. McAnally, X. Chen, and E.T. Kavalali et al. 2016. A peptide encoded by a transcript annotated as long noncoding RNA enhances SERCA activity in muscle. Science 351:271–275.

Olexiouk, V. W. Van Criekinge and G. Menschaert. 2018. An update on sORFs.org: a repository of small ORFs identified by ribosome profiling. Nucleic Acids Res 46:D497–D502.

Olexiouk, V., J. Crappe, S. Verbruggen, K. Verhegen, L. Martens, and G. Menschaert. 2016. sORFs.org: a repository of small ORFs identified by ribosome profiling. Nucleic Acids Res 44:D324–D329.

Oliveira, C.C. and J.E. McCarthy. 1995. The relationship between eukaryotic translation and mRNA stability. A short upstream open reading frame strongly inhibits translational initiation and greatly accelerates mRNA degradation in the yeast Saccharomyces cerevisiae. J Biol Chem 270:8936–8943.

Omasits, U., A.R. Varadarajan, M. Schmid, S. Goetze, D. Melidis, M. Bourqui, O. Nikolayeva, M. Quebatte, A. Patrignani, and C. Dehio et al. 2017. An integrative strategy to identify the entire protein coding potential of prokaryotic genomes by proteogenomics. Genome Res 27:2083–2095.

Pauli, A., M.L. Norris, E. Valen, G.L. Chew, J.A. Gagnon, S. Zimmerman, A. Mitchell, J. Ma, J. Dubrulle, and D. Reyon et al. 2014. Toddler: an embryonic signal that promotes cell movement via Apelin receptors. Science 343:1248636.

Peabody, D.S. 1989. Translation initiation at non-AUG triplets in mammalian cells. J Biol Chem 264: 5031–5035.

Pohl, M. G. Theissen and S. Schuster. 2012. GC content dependency of open reading frame prediction via stop codon frequencies. Gene 511:441–446.

Slavoff, S.A., A.J. Mitchell, A.G. Schwaid, M.N. Cabili, J. Ma, J.Z. Levin, A.D. Karger, B.A. Budnik, J.L. Rinn, and A. Saghatelian. 2013. Peptidomic discovery of short open reading frame-encoded peptides in human cells. Nat Chem Biol 9:59–64.

Somers, J. T. Poyry and A.E. Willis. 2013. A perspective on mammalian upstream open reading frame function. Int J Biochem Cell Biol 45:1690–1700.

Starck, S.R., V. Jiang, M. Pavon-Eternod, S. Prasad, B. McCarthy, T. Pan, and N. Shastri. 2012. Leucine-tRNA initiates at CUG start codons for protein synthesis and presentation by MHC class I. Science 336:1719–1723.

Sun, K., X. Chen, P. Jiang, X. Song, H. Wang, and H. Sun. 2013. iSeeRNA: identification of long intergenic non-coding RNA transcripts from transcriptome sequencing data. BMC Genomics 14 Suppl 2:S7.

Sun, L., H. Liu, L. Zhang, and J. Meng. 2015. lncRScan-SVM: A Tool for Predicting Long Non-Coding RNAs Using Support Vector Machine. PLoS One 10:e139654.

Tong, X. and S. Liu. 2019. CPPred: coding potential prediction based on the global description of RNA sequence. Nucleic Acids Res 47:e43.

Vilela, C. and J.E. McCarthy. 2003. Regulation of fungal gene expression via short open reading frames in the mRNA 5’untranslated region. Mol Microbiol 49:859–867.

von Arnim, A.G. Q. Jia and J.N. Vaughn. 2014. Regulation of plant translation by upstream open reading frames. Plant Sci 214:1–12.

Wang, G., H. Yin, B. Li, C. Yu, F. Wang, X. Xu, J. Cao, Y. Bao, L. Wang, and A.A. Abbasi et al. 2019. Characterization and identification of long non-coding RNAs based on feature relationship. Bioinformatics 35:2949–2956.

Wang, J., S. Li, Y. Zhang, H. Zheng, Z. Xu, J. Ye, J. Yu, and G.K. Wong. 2003. Vertebrate gene predictions and the problem of large genes. Nat Rev Genet 4:741–749.

Wang, L., H.J. Park, S. Dasari, S. Wang, J.P. Kocher, and W. Li. 2013. CPAT: Coding-Potential Assessment Tool using an alignment-free logistic regression model. Nucleic Acids Res 41:e74.

Wen, J. K.A. Lease and J.C. Walker. 2004. DVL, a novel class of small polypeptides: overexpression alters Arabidopsis development. Plant J 37:668–677.

Wethmar, K., A. Barbosa-Silva, M.A. Andrade-Navarro, and A. Leutz. 2014. uORFdb--a comprehensive literature database on eukaryotic uORF biology. Nucleic Acids Res 42:D60–D67.

Wucher, V., F. Legeai, B. Hedan, G. Rizk, L. Lagoutte, T. Leeb, V. Jagannathan, E. Cadieu, A. David and H. Lohi et al. 2017. FEELnc: a tool for long non-coding RNA annotation and its application to the dog transcriptome. Nucleic Acids Res 45:e57.

Zheng, J., X. Zhang, X. Zhao, X. Tong, X. Hong, J. Xie, and S. Liu. 2018. Deep-RBPPred: Predicting RNA binding proteins in the proteome scale based on deep learning. Sci Rep 8:15264.

Zhu, M. and M. Gribskov. 2019. MiPepid: MicroPeptide identification tool using machine learning. BMC Bioinformatics 20:559.

Zhu, Y., L.M. Orre, H.J. Johansson, M. Huss, J. Boekel, M. Vesterlund, A. Fernandez-Woodbridge, R. Branca, and J. Lehtio. 2018. Publisher Correction: Discovery of coding regions in the human genome by integrated proteogenomics analysis workflow. Nat Commun 9:1852.

